# Introducing risk inequality metrics in tuberculosis policy development

**DOI:** 10.1101/380865

**Authors:** M. Gabriela M. Gomes, Juliane F. Oliveira, Adelmo Bertolde, Tuan Anh Nguyen, Ethel L. Maciel, Raquel Duarte, Binh Hoa Nguyen, Priya B. Shete, Christian Lienhardt

**Author notes:** Correspondence and requests for materials should be addresses to M.G.M.G.

## Abstract

Global stakeholders including the World Health Organization rely on predictive models for developing strategies and setting targets for tuberculosis care and control programs. Failure to account for variation in individual risk leads to substantial biases that impair data interpretation and policy decisions^1,2^. Anticipated impediments to estimating heterogeneity for each parameter are discouraging despite considerable technical progress in recent years. Here we identify acquisition of infection as the single process where heterogeneity most fundamentally impacts model outputs, due to cohort selection imposed by dynamic forces of infection. Individuals with higher risk of acquiring infection are predominantly affected by the pathogen, leaving the unaffected pool with those whose intrinsic risk is lower. This causes susceptibility pools to attain average risks which are lower under higher forces of infection. Interventions that modify the force of infection change the strength of selection, and therefore alter average risks in the pools which feed further incidence. Inability to account for these dynamics is what makes homogenous models unsuitable. We introduce concrete metrics to approximate risk inequality in tuberculosis, demonstrate their utility in mathematical models, and pack the information into a risk inequality coefficient which can be calculated and reported by national tuberculosis programs for use in policy development and modeling.

---

Tuberculosis (TB) is a leading cause of morbidity and mortality worldwide, accounting for over 10 million new cases annually^4^. Although allusions are often made to the disproportionate effect of TB on the poorest and socially marginalized groups^5,6^, metrics to quantify inequality in risk to acquire TB are seldom used. Data reported by the World Health Organization (WHO), on which mathematical models often rely for calibrations and projections, are typically in the form of country-level averages.

Variation in individual characteristics has a generally recognized impact on the dynamics of populations, and pathogen transmission is no exception. In infectious diseases, the focus has been on heterogeneities in transmission through their nonlinear effects on the basic reproduction number, *R*_0_, in ways which are unique to these systems^7–11^. In TB, as in other communicable diseases, this approach motivated the proliferation of efforts to collect data on contact patterns and superspreading events, to unravel structures that may affect transmission indices and models. The need to account for variation in disease risk, however, is not unfamiliar in epidemiology at large, where so-called frailty terms are more generally included in linear models to improve the accuracy of data analysis^12^. The premise is that variation in risk of disease (whether infectious or not) goes beyond what is captured by measured factors, and a distribution of unobserved heterogeneity can be inferred from incidence trends in a holistic manner. Such distributions are needed for eliminating biases in interpretations and predictions, and can be utilized in conjunction with more common reductionist approaches, which are required when there is desire to target interventions at individuals with specific characteristics.

Individual risk of infection or disease relates to a probability of responding in a certain way to a stimulus and, therefore, direct measurement would require the recording of many exposures. In TB, this is unfeasible due to the relatively low frequency of disease episodes, but may be approximated by sub-dividing the population in sufficiently large groups^13^, according to some risk similarity criterion, and recording occurrences in each group. Then incidence rates can be calculated per group, and ranked. Extended Data Fig. 1 illustrates the population of a hypothetical country comprising low and high risk individuals distributed geographically (but grouping by income level, for example, or a combination of the two could also serve our statistical purposes). Forasmuch as individuals are non-uniformly distributed, disease incidence will vary between divisions and carry information about intrinsic relative risks (Methods, and Extended Data Figs. 2 and 3).

Fig. 1 refers to the population of Brazil divided into municipalities. Incidences were ranked, a Lorenz curve^14^ was generated (Fig. 1a) and a high-incidence quantile characterized (4% was considered, but the procedure would apply for any desired partition), leading to a discretization into two incidence levels (Fig. 1d). A mathematical model for tuberculosis transmission with two risk groups^1,15^ was solved (Methods, Extended Data Fig. 4) and the relative risk parameters estimated by fitting the relative incidences calculated from the data available for 2002 (Fig. 1b, c). The procedure was applied to data from Vietnam, Brazil and Portugal (countries with high, medium and low TB burdens, respectively), providing variance estimates of 5.4 in Vietnam, 8.9 in Brazil and 0.84 in Portugal. Notwithstanding the conservative nature of risk variances estimated by this procedure, in-depth comparative analysis reveals the mechanism behind the poor predictive capacity of homogeneous models and identifies acquisition of infection as the single most important process in this phenomenon.

**Fig. 1:**
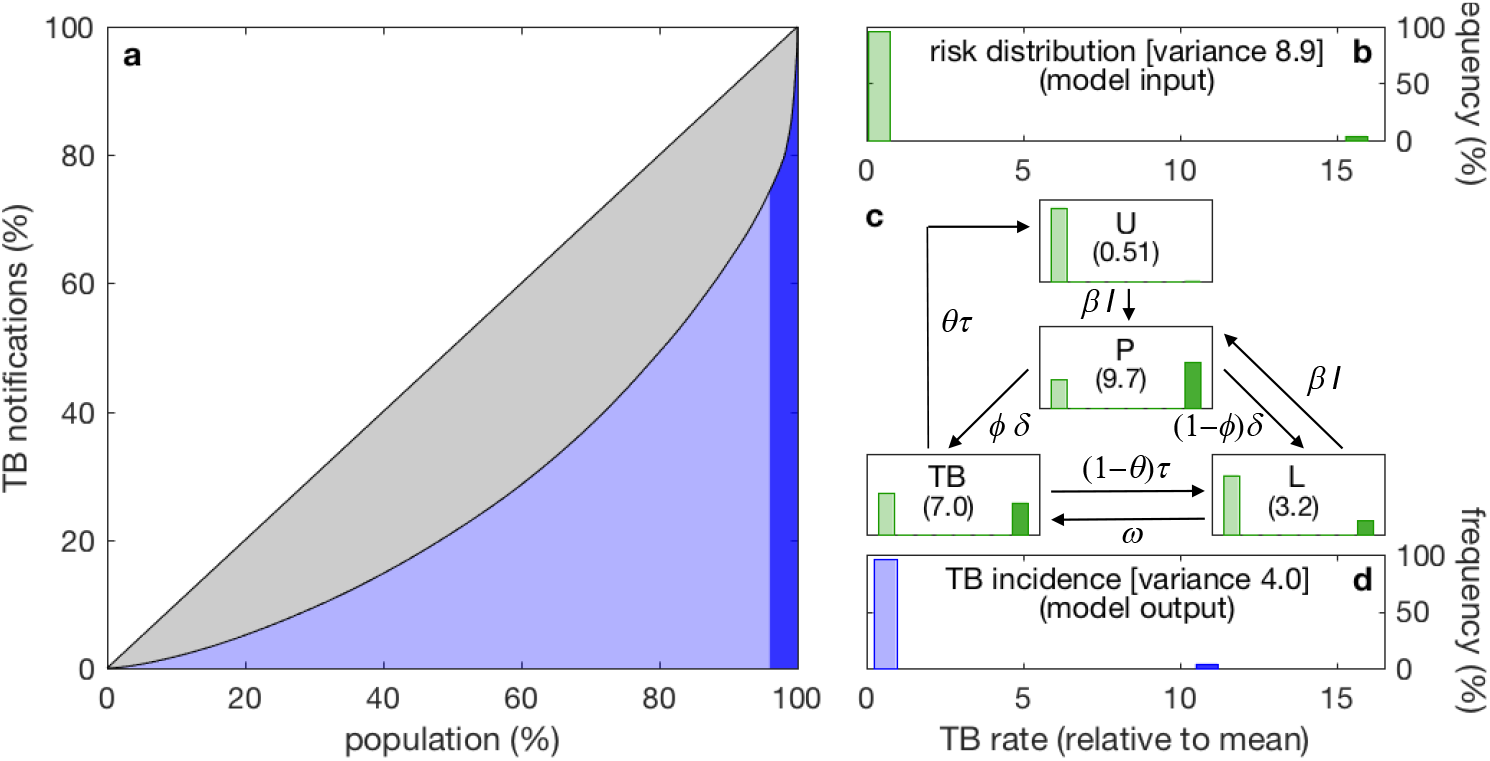
Risk distributions and transmission model. **a**, Lorenz curve^14^ constructed by ordering municipalities in a country from lower to higher TB notification rates (Brazil adopted for illustration), for calculating the notification rate in the 4% higher-risk quantile (darker blue) in relation to the population average. **b**, Risk distribution required by a mathematical model (Methods) for generating relative incidence rates compatible with the 2002 notification data (*α*_1_ = 0.53, *β*_2_ = 12 in Vietnam [variance 5.4]; *β*_1_ = 0.39, *α*_2_ = 16 in Brazil [variance 8.9]; *α*_1_ = 0.81, *β*_2_ = 5.5 in Portugal [variance 0.84]). **c**, Risk distributions in the various epidemiological compartments segregated by the transmission dynamics calibrated by the 2002 incidence for Brazil (52 per 100,000 person-years, as per WHO’s global tuberculosis database), **d**, Distribution of incidence rates discretized into two risk groups (*I*_1_ = 0.80, *I*_2_ = 5.8 in Vietnam [variance 0.96]; = 0.59, *I*_2_ = 11 in Brazil [variance 4.0]; = 0.88, *I*_2_ = 4.0 in Portugal [variance 0.37]). Model parameters (Extended Data Table 1): *β* = 3.47 *yr*^−1^; *ϕ* = 0.05; *δ* = 2 *yr*^−1^; *θ* = 1; *τ* = 2 *yr*^−1^; *ω* = 0.0013 *yr*^−1^; *μ* = 1/80 *yr*^−1^. Notice that observed incidence variances (or ratios) invariably indicate underlying risk variances which are higher^12^.

Extended Data Fig. 5 depicts the action of cohort selection in TB epidemiology. The model fitting the incidence of Brazil in 2002 was solved numerically with a constant decline in reactivation rate to meet an arbitrary fixed target of halving the incidence in 10 years (Extended Data Fig. 5a, black curve). If these estimations and projections had been made by the mean field approximation of the same model, the required control efforts would have been underestimated and the target systematically missed (Methods, and Extended Data Table 2). This is because the force of infection decreases as the intervention progresses, relieving the selection pressure on epidemiological compartments at risk of infection, which consequently undergo an increase in mean risk, opposing the intended effects of the intervention (Extended Data Fig. 5b). Homogeneous models artificially disable cohort selection, creating an illusion that control targets are moving when observed from a homogeneous frame. This is a general phenomenon in infectious diseases, but we find that in TB the sign of the effect (i.e. whether homogeneous models *overestimate* or *underestimate* impact) is not the same for every intervention due to a buffering effect exerted by the latent compartment.

More practically, the model was used to reproduce reported country-level trends for TB incidence in Vietnam, Brazil, and Portugal. The model was initialized in 2002 as before, but the incidence declines reported by WHO until 2015 were shared equally among three relevant parameters (Methods): probability of progression from primary infection to active disease (*ϕ*, with the remaining 1 − *ϕ* maintaining a latent infection); rate of reactivation of latent infection (*ω*); rate of successful treatment (*τ*). Reported annual incidences and model trajectories prolonged until 2020 are shown in fig. 2a, e, i, and the proportional improvements in each of the three varying parameters are shown in fig. 2c, g, k.

**Fig. 2:**
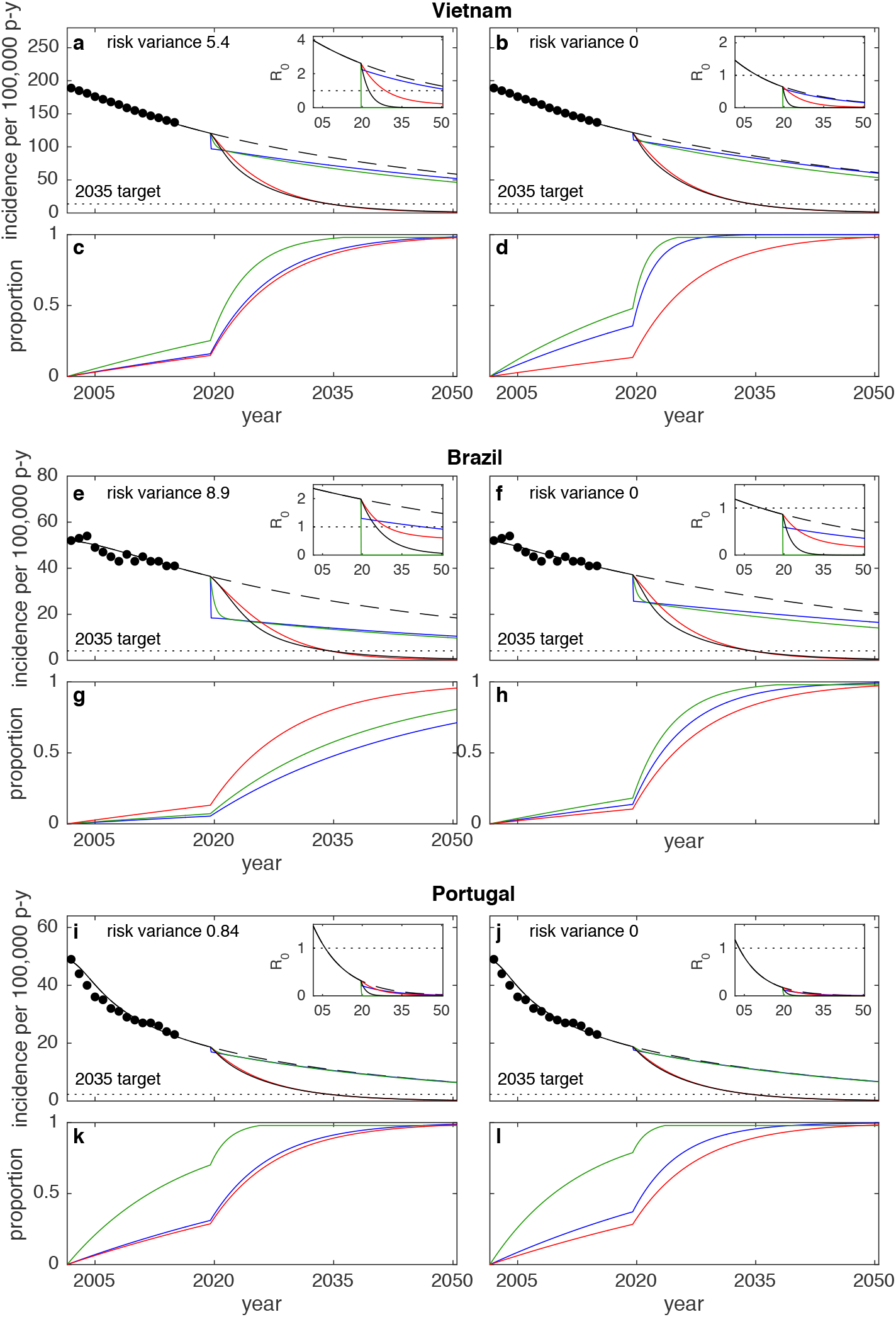
Heterogeneous vs homogeneous model. TB incidence from 2002 to 2015 (black dots), and model solutions under heterogeneous (**a**, **e**, **i**) and homogeneous (**b**, **f**, **j**) infection risks. Initial parameters values calculated by adjusting effective contact rates (β) to fit 2002 incidence rates, and declines towards 2015 incidences (solid black curve) shared in equal parts between proportion progressing from primary infection to active disease (f), rate of reactivation of latent infection (*ω*), and cure rate (*τ*) (Methods, Extended Data Table 3). Other parameters as in Extended Data Table 1, with 0 = 1. From 2020 onwards, the trajectories split to represent four scenarios: rates of parameter change are maintained (dashed black); scale change in f, *ω*, and *τ*to meet WHO target for 2035 (black); scale change in *f* only (blue), **ω** only (red), or *τ*only (green). (**c, d**) Proportional improvements in *f* (blue), *ω* (red) and **τ** (green) associated with the solid black trajectories of (a, b). (**g**, **h**) and (**k**, **l**) Same in relation to (e, f) and (i, j), respectively. The main top panels for each country are very similar because they were generated by fitting two model versions to the same data, but with evidently different underlying parameter values (top panel insets and bottom panels).

Incidence targets set by the WHO for 2035 are marked in fig. 2a, e, i by the dotted lines, and the model projections for continuing the same parameter decline rates beyond 2020 are represented by the dashed curve segments. Evidently, control measures must be intensified for meeting the ambitious targets. For each country, we apply a uniform scaling factor (*κ*) to the three parameters which is calculated to represent the required efforts (*κ* = 13 in Vietnam; *κ* = 12 in Brazil; *κ* = 6 in Portugal) (Methods, Extended Data Table 4). A homogeneous analogue of the model would adjust the data undistinguishably well (Fig. 2b, f, j), but require different efforts in parameter changes (Fig. 2d, h, l) specially pertaining to treatment rate (*τ*) and progression probability (f). Specifically, the homogeneous model would overestimate required changes in treatment success and progression from primary infection to active disease, implying that active detection of cases and respective contacts appears relatively more rewarding under heterogeneity (contrary to interventions that emphasize reduction in reactivation rates, which appear more impactful under homogeneity^1^). This is attributed to an inflation of the relative contribution of reactivation in homogeneous settings which also results from the weakening of cohort selection as follows.

High risk individuals are disproportionately affected by the pathogen, leaving behind unaffected pools with reduced mean susceptibilities. The strength of this selection effect is determined by the interplay between risk variance and the force of infection, appearing stronger in Brazil than in Vietnam, and weakest in Portugal (Fig. 3). Homogeneous models disable this natural process and, thereby, promote an artificial impression of low transmission parameters^16^ (such as *β* or *R*_0_, compare insets in fig. 2 a-b, e-f, i-j). System feedbacks compensate this underscoring in transmission by inflating the role of reactivation to meet reported incidences and therefore reducing the impact basis for interventions focused on ongoing transmission. Overall, this suggests that current emphasis on active case finding and contact investigation^17–19^ is likely to be more effective than homogeneous models suggest.

**Fig. 3:**
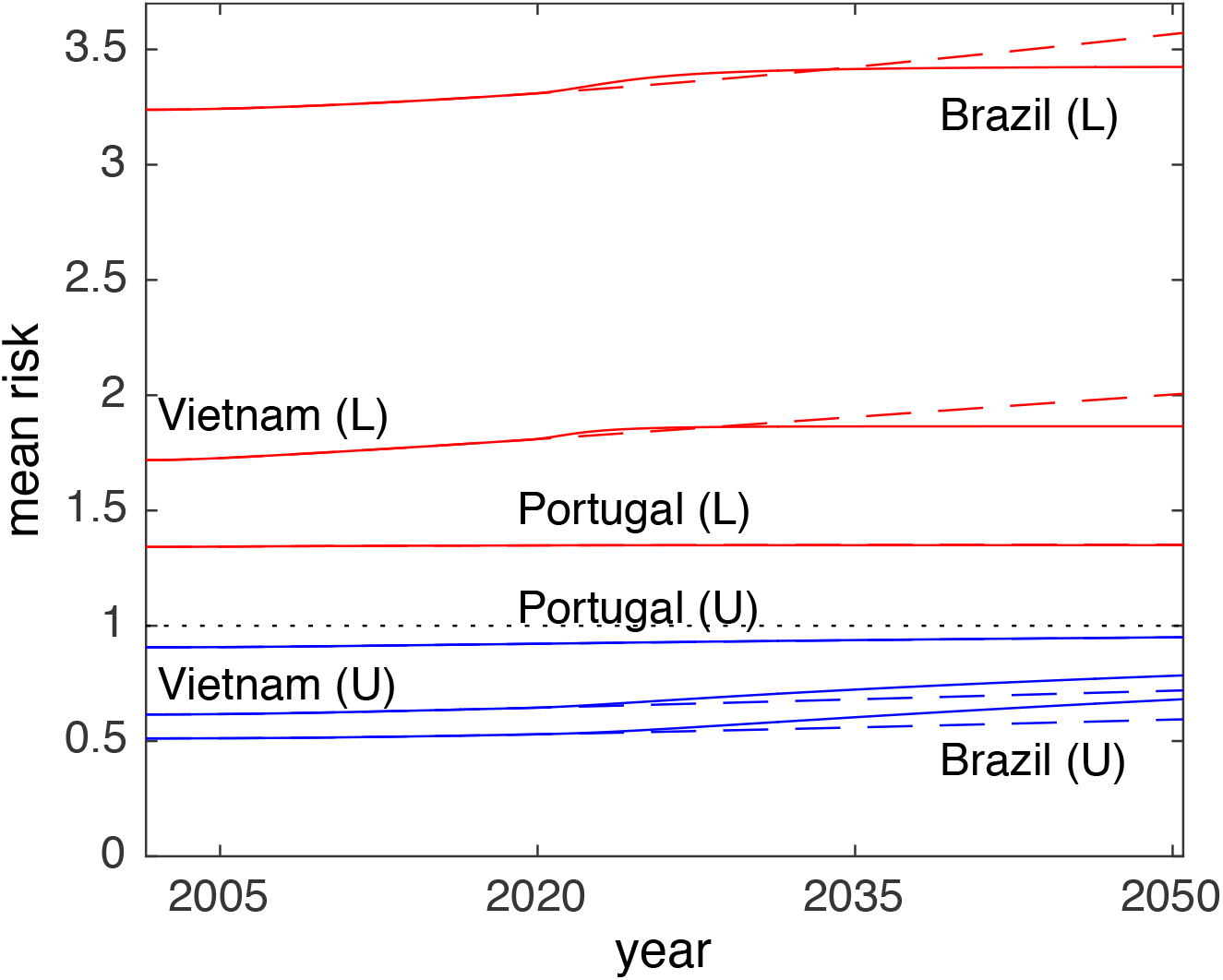
Risk distribution dynamics. Mean risk of infection in the latent and uninfected compartments (Methods). Solid and dashed curves associate, respectively, with solid and dashed black trajectories in fig. 2 a, e, i.

The colored trajectories in fig. 2a, e, i represent model solutions where the intensification of control efforts from 2020 onwards is focused on single parameters. The outcomes are consistently outperformed by integrated efforts that address all three parameters simultaneously (in black, as above) and, specifically, addressing reactivation is crucial for meeting the targets in all scenarios. Prevalences of latent TB infection (LTBI) calculated from our model trajectories (30.8% in Vietnam, 16.6% in Brazil, and 19.7% in Portugal, in 2014) are consistent with estimates from a recent study^20^. Even though these percentages are somewhat smaller than those expected by the homogeneous model counterpart (Extended Data Table 5), and heterogeneity reduces the impact of interventions that primarily reduce reactivation rates^1^ (Extended Data Fig. 5), the reservoir must nevertheless be reduced if incidence targets are to be met. Extended Data Fig. 6 depicts cumulative reductions in reactivation rates associated with fig. 2, consistently indicating that reducing reactivation rates by about 80% (84-87% in Vietnam; 78-90% in Brazil; 84-85% in Portugal, where lower bounds correspond to integrated interventions and upper bounds result from reducing reactivation only) is a requirement for meeting the End TB incidence targets by 2035.

The variation in risk addressed here considers that acquisition of infection is positively correlated with transmission, as expected when heterogeneity is due to contact patterns^1,7,9–10,21^. In the Supplementary Information, alternative assumptions are examined. First, we show that the results are robust to whether individuals clear the infection upon treatment or maintain latent infection (Supplementary figs. 1-3), and whether acquisition of infection is correlated with transmission or not (i.e. whether variation is in degree of contacts or susceptibility to infection (Supplementary fig. 4)). Second, we show that when variation is in the probability of progression from primary infection to active disease, model outputs do not deviate from the homogeneous model because progression is not under the cohort selection mechanisms described above (conveyed by the trajectories of mean risk not moving away from one in supplementary fig. 5c, e, g). Similarly, variation in rates of reactivation or treatment success should not lead to different model outputs unless correlated with propensities for acquiring infection. Third, we show that halving susceptibility due to acquired immunity from exposure to the pathogen does not change the direction of our conclusions (Supplementary figs. 6-8), although this may increase the size of the reported effects in the highest incidence regions due to the induction of a reinfection threshold^1,22^.

The notion that heterogeneity affects the results of population models and analyses is not new^3,23–25^, but we still face a general inability to measure it. We advance a concrete way forward for infectious disease transmission models, which is based on routinely collected data. Measures of statistical dispersion (such as the quantiles used here or the Gini coefficient^26^) are commonly used in economics to represent the distribution of wealth among individuals in a country, and to compare inequality between countries, but rarely used in epidemiology^27^. Measuring disease risk of an individual is less direct than measuring income, but surely this can be overcome in creative ways for specific diseases.

We have focused on tuberculosis, and shown how to approximate distributions of individual risk from suitably structured disease notification and population data (Fig. 4), and how to summarize the information into a simple risk inequality coefficient (*RIC* = 0.30 in Vietnam, *RIC* = 0.46 in Brazil, and *RIC* = 0.32 in Portugal), analogous to the Gini coefficients estimated by the World Bank (0.38 in Vietnam, 0.51 in Brazil, and 0.36 in Portugal). We have demonstrated how to feed this information into tractable mathematical models and why this is essential to predictive capacity. The worldwide adoption of such an index would enable policymakers and key public health programs to assess risk inequality in each country, compare the metric across countries, and monitor the impact of equalization strategies and targeted interventions over time.

**Fig. 4:**
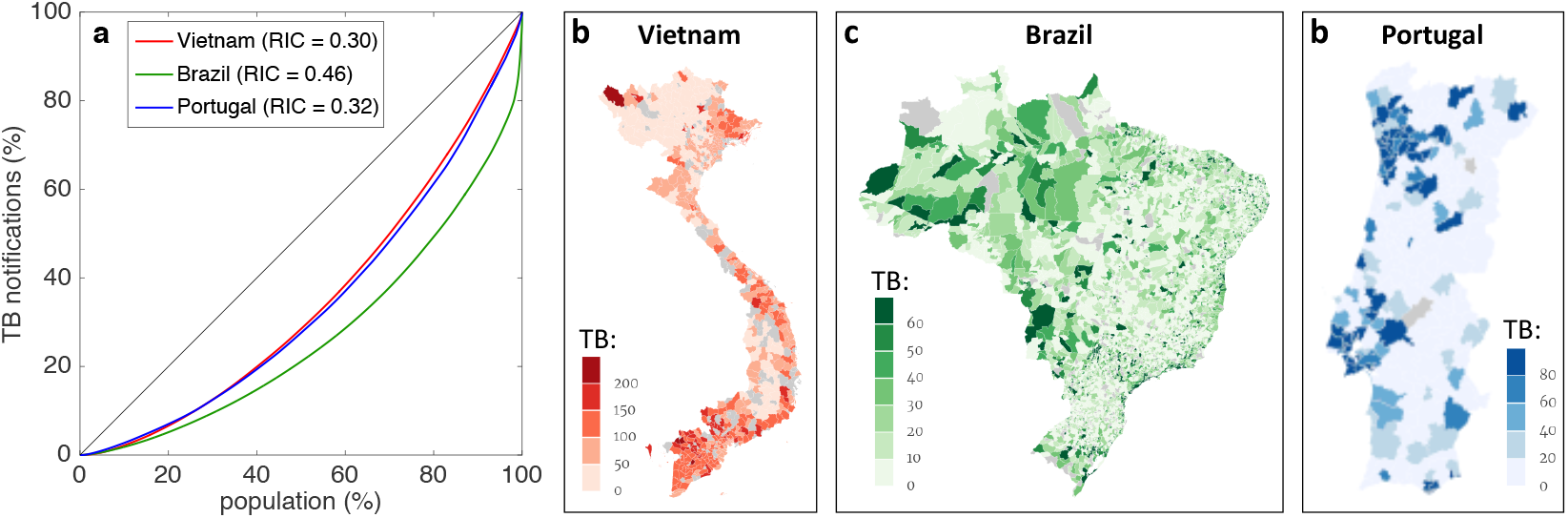
Risk inequality coefficient. **(a)** Lorenz curves calculated from notification data stratified by level 2 administrative divisions (697 districts in Vietnam; 5127 municipalities in Brazil; 308 in Portugal). Risk inequality coefficients (RIC) were calculated from Lorenz curves as in Methods and Extended Data Fig. 3. Country maps with administrative divisions for Vietnam **(b)**, Brazil **(c)**, and Portugal **(d)**, colored by number of cases notified by 100,000 person-years.

## METHODS

### Lorenz curves and risk inequality coefficients

Lorenz curves^14^ are widely used in economics to calculate indices of inequality in the distribution of wealth, known as Gini coefficients^26^. Although rarely used in epidemiology, similar metrics can be used to describe inequalities in disease risk^27^. Here we construct a Lorenz-like curve for each study country from TB notifications and population data structured by municipalities (level 2 administrative divisions). Municipalities are ordered by incidence rates (from low to high) and cumulative percentage TB notifications are plotted against cumulative percentage population. By construction, this results in a convex curve between (0,0) and (100,100), which would be a straight line in the absence of inequality. A risk inequality coefficient (RIC) can be calculated as the ratio of the area between the curve and the equality line, over the area of the triangle under the equality line (Extended Data Fig. 2). This gives a number between 0 and 1, which is analogous to the Gini coefficient used to describe income inequality, with the exception that while income can be measured at the individual level for the constructions of Lorenz curves, the assessment of TB risk cannot be made by analyzing individuals directly, but must be approximated from group measurements.

In this paper, we use Lorenz curves to inform risk distributions for TB transmission models and propose their use more widely. However, this requires access to detailed datasets, which may be impractical at the global level. This incommodity may be circumvented by having National Tuberculosis Programs (NTP) to pack the information into a RIC summary that modelers can then unpack as appropriate for their models.

Extended Data Fig. 3 compares alternative Lorenz curves generated for Vietnam, Brazil and Portugal to explore the effects of timespan and group size. As we must comply with the administrative divisions already established in each country, level 2 appear to offer the best compromise between resolution (the smaller the units, the closer we get to measuring individual risk) and occurrences (the larger the units, the larger the numbers and the better the risk discrimination^13^). Regarding timespan, the longer the data series the better.

### Mathematical model

We adopt a TB transmission model adapted from previously published studies^1,15^,

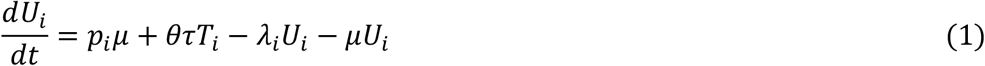

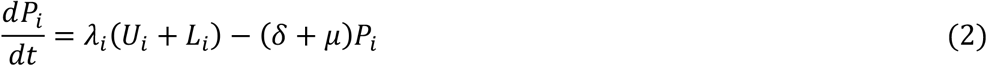

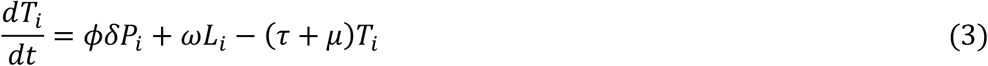

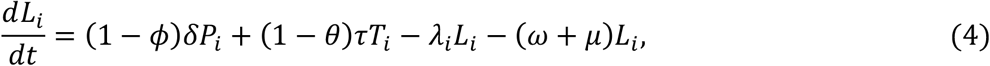

where subscripts *i* = 1,2 denote low and high risk groups, and within each group individuals are classified, according to their infection history, into uninfected (*U_i_*), or infected in one of three possible states: primary infection (*P_i_*); latent infection (*L_i_*); active tuberculosis disease (*T_i_*). The model parameters along with their typical values used herein are listed in Extended Data Table 1. The force of infection upon uninfected individuals is

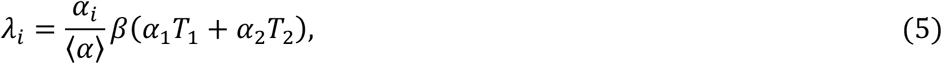

where *a_i_* is the “risk” of individuals in group *i* in relation to the population mean 〈*α*〉 = 1, and the basic reproduction number is

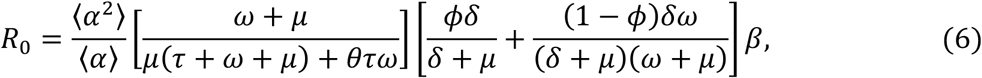

where 〈*α*^2^〉 is the second moment of the “risk distribution”. We use “risk” and “risk distribution” as generic terms to designate factors of variation in the propensity of individuals to acquire infection, which may be realized physically as intensity of exposure (Main Text) or biologically as susceptibility to infection given exposure (Supplementary Information). We use the terminology “epidemiological compartment” to refer to the composite of all compartments for the same infection status (i.e. “uninfected” comprises both *U*_1_ and *U*_2_, etc). We also introduce the notion of “mean risk” for each epidemiological compartment to track cohort selection (e.g. the mean risk for *U*(*t*) is calculated as *U*_1_+*U*_2_(*t*)*α*_2_, etc). We adopt two risk groups for concreteness, but any formalism for two or more groups would essentially accommodate the same phenomena. Indeed, two recent studies implemented cohort selection in populations structured into hundreds of risk groups^28,29^.

The model admits an endemic equilibrium when *R*_0_ > 1, as displayed by the solution curves parameterized by *β* in Extended Data Fig. 4. Incidence rates in each risk group are calculated from model outputs by the formula *I_i_* = *ϕδP_i_* + *ωL_i_* per year, and for the entire population as the sum of these over risk groups.

### Model initialization

Model trajectories are initialized assuming equilibrium conditions in 2002. Parameters describing the rates of birth and death of the population, the probability of progression from primary infection to active disease, and the rate of successful treatment, are set at the same values for the three countries: *μ* = 1/80 *yr*^−1^; *ϕ* = 0.05 (Ref. 30), *τ* = 2 *yr*^−1^(Ref. 31). The rate of reactivation is considered three times higher in South East Asian than in Western populations: *ω* = 0.0013 *yr*^−1^ in Brazil and Portugal; *ω* = 0.0039 *yr*^−1^ in Vietnam (Ref. 32). The mean effective contact rate (*β*) was calibrated to enable model solutions to meet country-level incidences estimated by the WHO for 2002 (Extended Data Fig. 4). Risk group frequencies are set at *p*_1_ = 0.96, and *p*_2_ = 0.04, and the relative risk parameters (*α*_1_ and *β*_2_) estimated as described below. The results are then displayed in terms of the non-dimensional parameter *R*_0_ (Extended Data Fig. 4b), which is linearly related to *β* according to (6).

The same procedure was carried out for the mean field approximation of the same model (Extended Data Fig. 4b). At this point it can be noted that *R*_0_ estimates are consistently higher under homogeneity^1^.

### Risk distributions

Given a Lorenz curve (Fig 1a), any quantile can be selected to characterize concentration of risk. We adopt “4% of the population account for *Y*% of the TB cases”, but the conclusions are not specific to the chosen quantile. A distribution of incidences is thus constructed directly from the data, such as: a segment *p*_1_ = 0.96 of the population accounts for (100 − *Y*)% of the incidence, while the remaining segment *p*_2_ = 0.04 accounts for the remaining *Y*% (Fig 1d). Model (1-5) is then solved as above, and the relative risk parameters *α_i_* are calculated (Fig. 1b) so as to output the desired incidence distributions. This was performed numerically by binary search to adjust the variance in the parameters *α_i_* such that the variance in the output incidences agrees with the notification data.

Under any positive force of infection, the two risk groups segregate differently to populate the various epidemiological compartments, as depicted in fig. 1c, resulting in mean risks that differ from 1 for specific compartments, and thereby deviating from homogeneous approximations. Crucially, the mean risks among individuals that occupy the various epidemiological compartments (brackets in fig. 1c for the conditions of Brazil in 2002) respond to dynamic forces of infection causing divergence from predictions made by homogenous models.

### Moving target

The model, with the estimated risk distribution, parameters, and initial conditions, fitting the incidence of Brazil in 2002 (52.0 per 100,000), is solved forward in time with a constant decline in reactivation rate as to meet an arbitrarily fixed target of halving the incidence in 10 years (to achieve 26.0 per 100,000). As in the calculation of risk variance above, also here we refer to a simple numerical calculation performed by binary search. We write the reactivation rate as *ω*(*t*) = 0.0013*e*^*r_ω_*(*t* − 2002)^ per year, and approximate *r_ω_* in order to meet the assumed incidence target by year 2012.

So, starting with an initial reactivation rate of 0.0013 per year, we find that meeting the target by this strategy alone, would require *r_ω_* = −0.1087, or equivalently a decline in reactivation by 1 − *e^r_ω_^* = 10% each year. This is to say that, in 10 years, the reactivation rate would have been reduced to approximately 0.0004 (heterogeneous set of columns in Extended Data Table 2).

Suppose that these estimations and projections were being made by the mean field approximation of the same model, and the outcomes were monitored yearly and readjusted if necessary. The result would have been that *r_ω_* = −0.0732 would suffice, corresponding to a yearly decline in reactivation rates of only 1 − *e^r_ω_^* = 7% to meet the target. Since the real population is heterogeneous, however, we simulate this decline for the first year with the heterogeneous model. The result is that, instead of achieving the projected incidence of 49.6 per 100,000, the measure lags with a reduction to only 50.4 per 100,000, a result that the homogeneous model would attribute to a failure in the effort to reduce reactivation. Indeed, from the homogeneous frame, an observer would have erroneously concluded that the decline had been by only *r_ω_* = −0.0473, and would have re-estimated the effort to meet the target over the remaining 9 years, now with an intensification to compensate for the lag of the first year (*r_ω_* = −0.0765). This process is simulated recursively for 10 years to populate Extended Data Table 2 and generate Extended Data Fig. 5. The inset in Extended Data Fig. 5a depicts the relative error committed each year.

The dynamics of the mean risk of infection in the uninfected and latent compartments as the described interventions proceeds are shown in Extended Data Fig. 5b to demonstrate the action of cohort selection. This is the key process leading to the deviation between the homogeneous and heterogeneous models.

### Control measures

The model with initial conditions, parameters, and distributions estimated in 2002, is used to reproduce reported country-level trends for TB incidence in Vietnam, Brazil, and Portugal. Incidence declines between 2002 and 2015, reported by WHO for each of the three countries, are assigned to changes in parameters *ϕ*, *ω*, and *τ*. The decline is shared equality among the three parameters, as a compromise, since identifiability of the precise share and estimation of respective uncertainty requires additional discriminatory data^33^, and is not essential for the purposes of this paper. This approach leads to exact solutions which can be approximated numerically using binary search algorithms.

As interventions proceed we monitor the improvements being made on the target parameters. Improvements correspond to decline in the probability of progression from primary infection to active disease [1 − *ϕ*(*t*)/*ϕ*(2002)], decline in reactivation rate [1 − *ω*(*t*)/*ω*(2002)], or increase in the rate of successful treatment [1 − *τ*(2002)/*τ*(*t*)].

## Acknowledgements

The Bill and Melinda Gates Foundation is acknowledged for its support through grant project number OPP1131404. MGMG and JFO received additional support from Fundação para a Ciência e a Tecnologia (IF/01346/2014).

## Author contributions

M.G.M.G., P.B.S, and C.L. designed the study; E.L.M., R.D., and B.H.N. provided NTP data and expertise; M.G.M.G., J.F.O., A.B., and T.A.N. performed the analysis; M.G.M.G., P.B.S, and C.L. drafted the manuscript; all authors revised and approved the final version.

## Data availability

Estimated country-level incidence rates obtained from the WHO’s global tuberculosis database (http://www.who.int/tb/country/data/download/em/), and municipality-level notification and population data from the respective National Tuberculosis Programs.

## Supplementary Information

This file contains Supplementary Text and Supplementary Figures 1-8.

**Extended Data Fig. 1:**
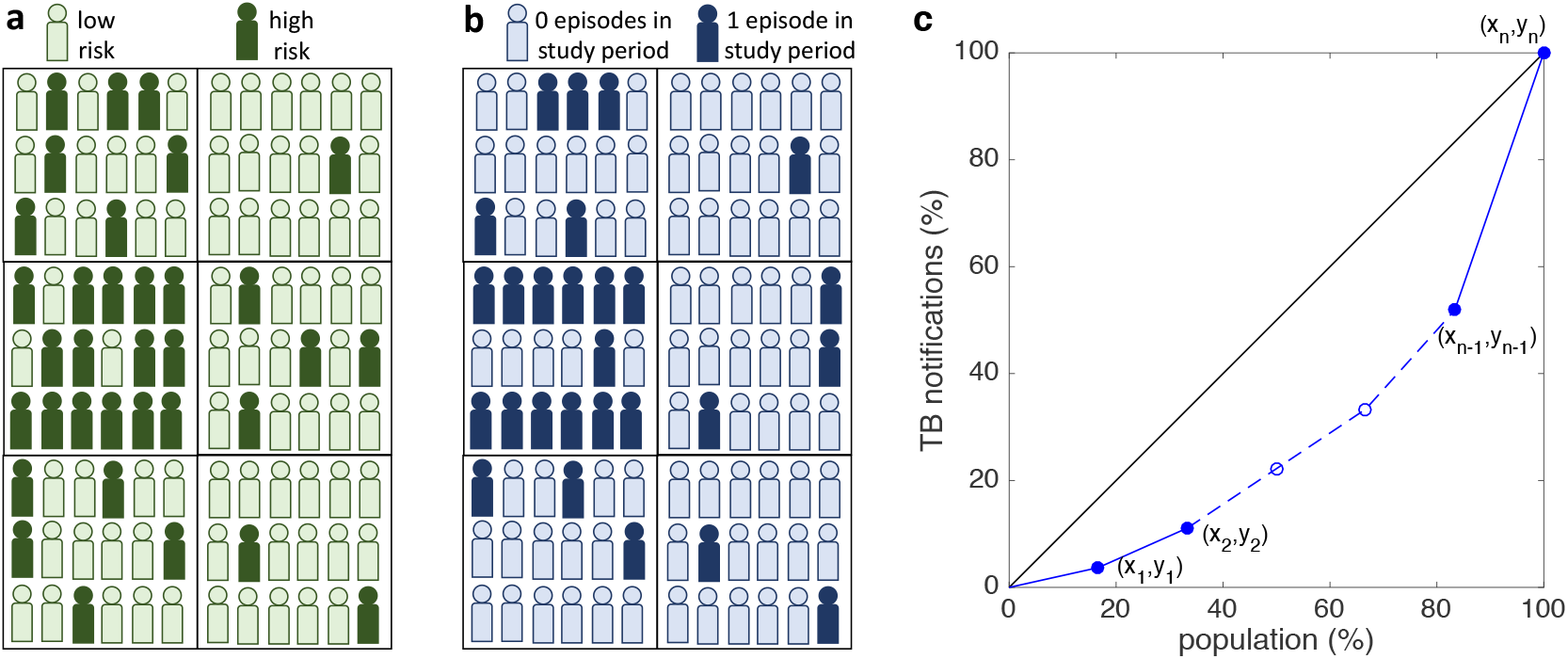
Illustration of a non-uniform distribution of low and high risk individuals. A non-uniform distribution of low and high risk individuals (**a**) leads to variation in disease incidence across population divisions (**b**), with divisions with higher frequency of high-risk individuals naturally having higher incidence rates. In reverse, conservative measures of risk inequality can be obtained from conveniently stratified incidence data: **c** depicts the construction of a Lorenz-like curve^14^ from incidence data in a hypothetical country partitioned into geographical divisions, for instance (b). Population divisions are ranked from lower to higher incidence and cumulative measures of population (*x_i_*) and disease notifications (*y_i_*) are calculated for every index *i* between 1 and the total number of divisions *n*. Lorenz curves are widely used in economics to summarize measures of inequality in the distribution of wealth and can be used in creative ways to describe distributions of disease risk in populations.

**Extended Data Fig. 2:**
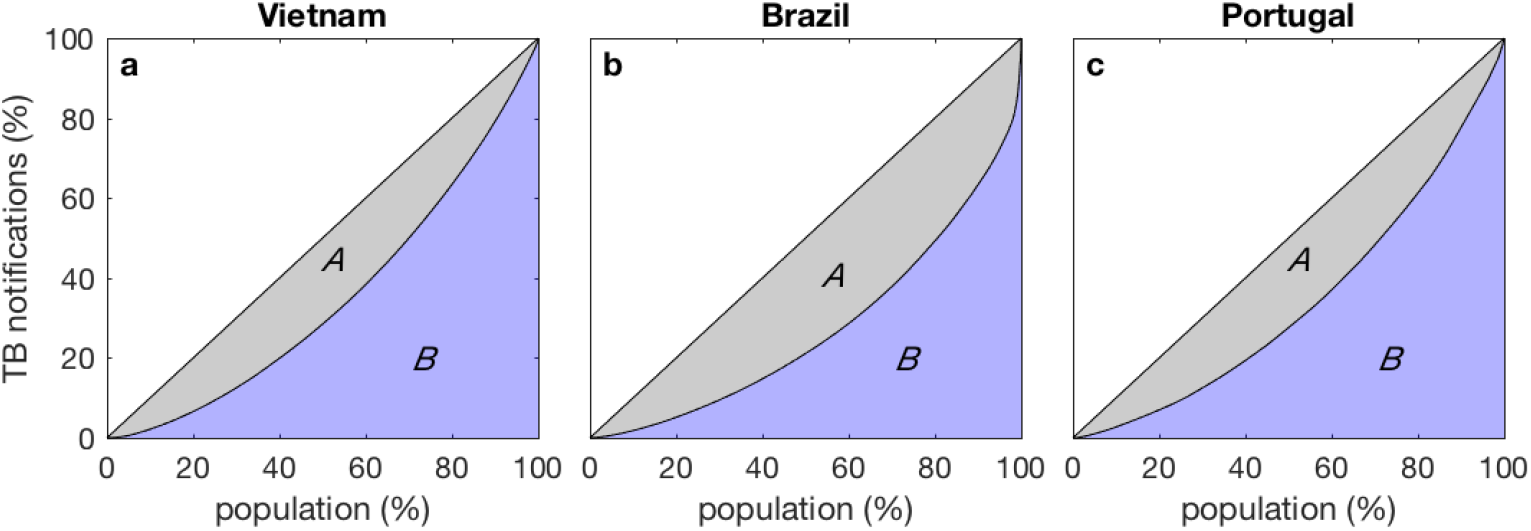
Risk inequality coefficient (RIC). Lorenz curves from TB notifications by municipality in Vietnam (**a**), Brazil (**b**), and Portugal (**c**). The risk inequality coefficient is calculated as *RIC* = *A*/(*A* + *B*). where *A* is the area between the Lorenz curve and the line of equality (grey), and *B* is the (blue) area under the Lorenz curve 04 +*B* is the total shaded area).

**Extended Data Fig. 3:**
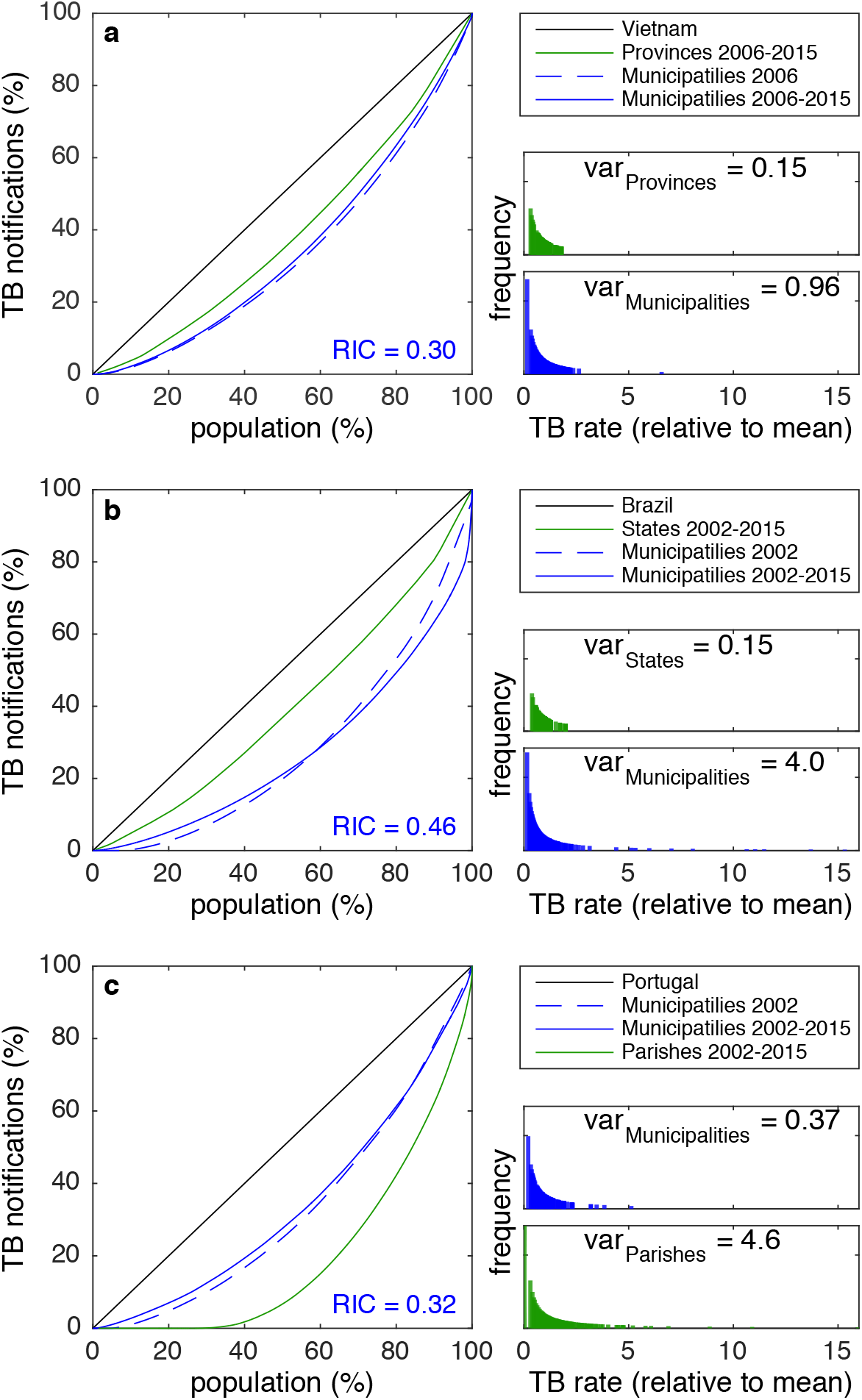
Risk inequality metrics. Lorenz curves calculated from notification data covering various time periods and administrative divisions. Level 2 divisions (**a** 697 municipalities in Vietnam; **b** 5127 in Brazil; **c** 308 in Portugal) are shown in blue (solid using the number of years available to this study [2006-2015 in Vietnam, and 2002-2015 in Brazil and Portugal], and dashed using only the first year in each series). Green curves correspond to alternative administrative levels (64 provinces in Vietnam [level 1]; 27 states in Brazil [level 1]; 3281 parishes in Portugal [level 3]). Notification rate distributions corresponding to blue and green solid curves are plotted together with the respective variances. Variances obtained from the blue distributions were used in models throughout this paper. Two-risk group discretizations were used in the paper while here we show 100-risk group discretizations of the same distributions. The entire analysis could be carried out with more resolved distributions if desired by simply increasing the dimension of the system accordingly.

**Extended Data Fig. 4:**
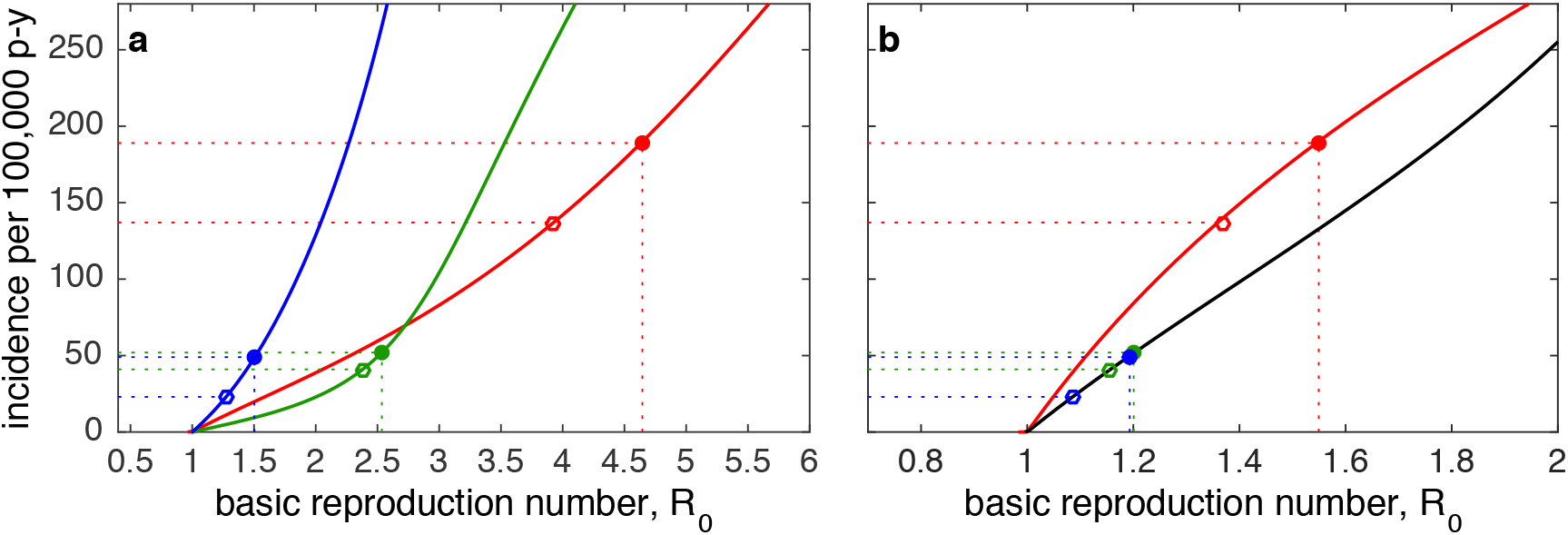
Endemic equilibria and incidence rates in 2002 and 2015. Equilibrium solutions of model (1-6) with risk distributions estimated for Vietnam (red), Brazil (green), and Portugal (blue) (**a**), and the mean field approximation of the same model (**b**). Parameter values: *ϕ* = 0.05; *ω* = 0.0039 *yr*^−1^ (Vietnam); *ω* = 0.0013 yr^−1^ (Brazil and Portugal); *τ* = 2 *yr*^−1^; *δ* = 2 *yr*^−1^; *μ* = 1/80 *yr*^−1^; *θ* = 1. Filled and open circles mark incidence rates in 2002 and 2015, respectively, as reported by WHO.

**Extended Data Fig. 5:**
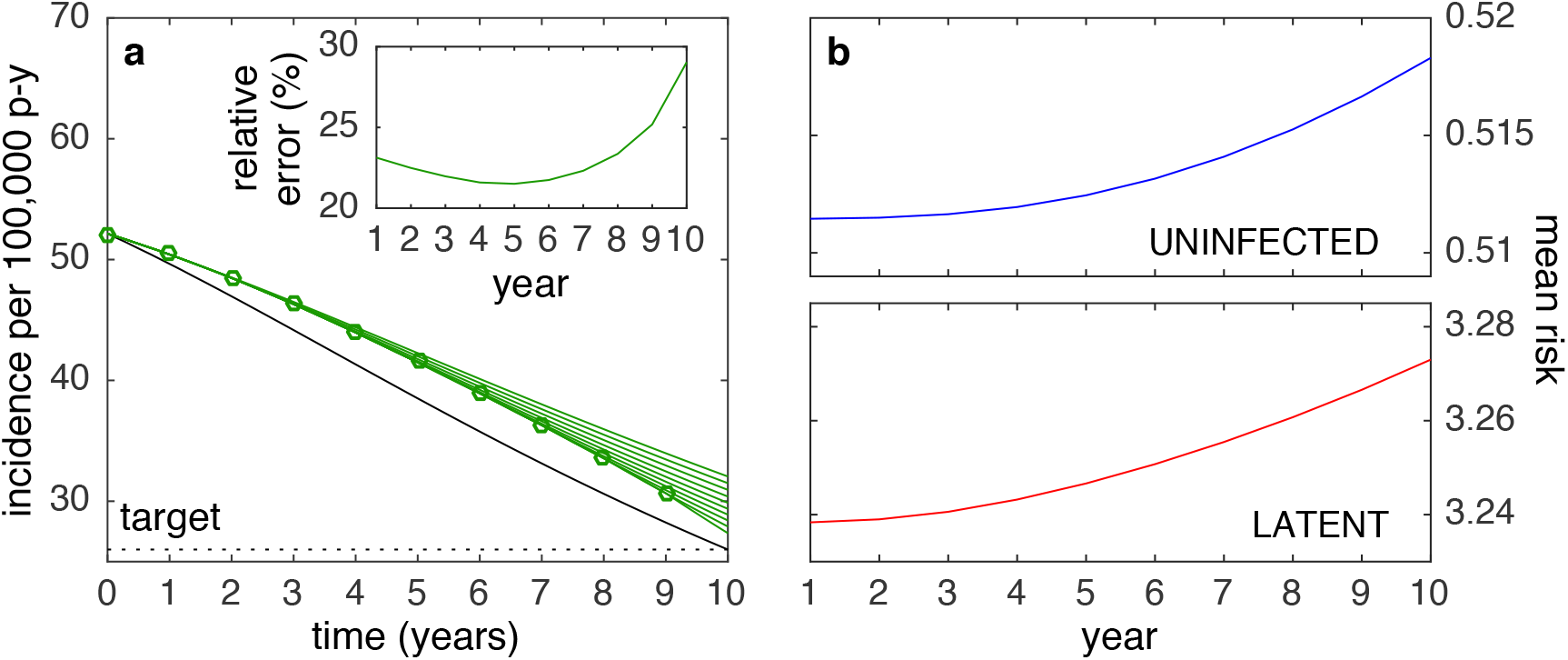
Moving target. How (**a**) and why (**b**) fixed targets appear to be moving when observed from a homogeneous frame (Methods, and Extended Data Table 2).

**Extended Data Fig. 6:**
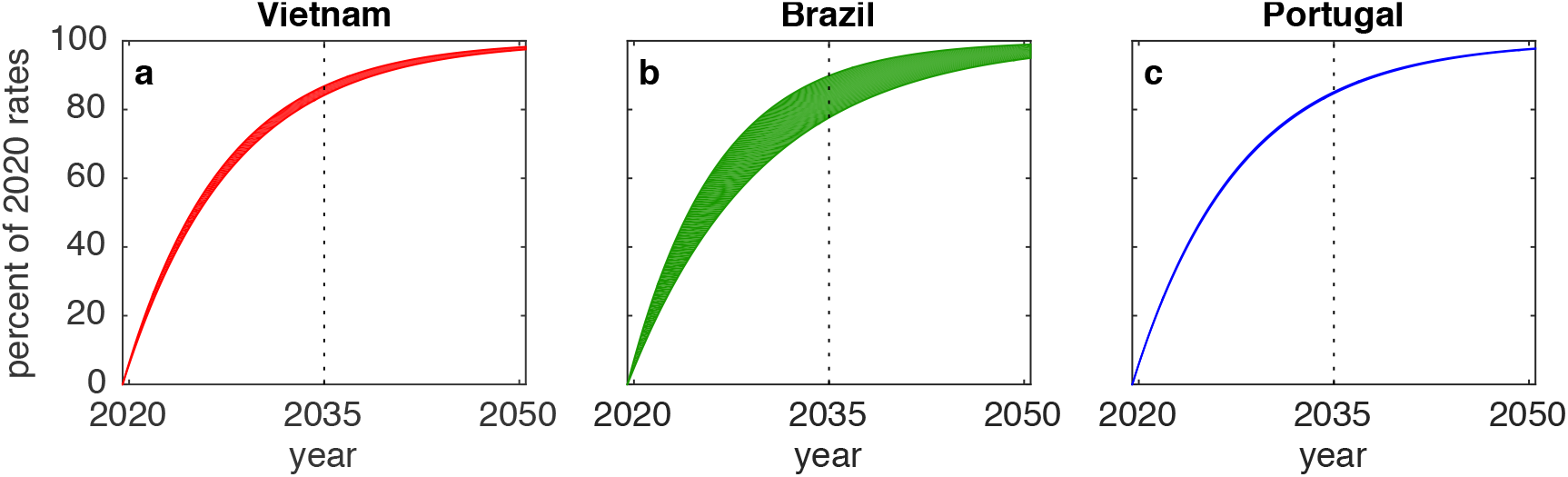
Cumulative reduction in reactivation rates required for meeting End TB incidence targets. Calculated by integrating annual declines obtained in fig. 2, for Vietnam (**a**), Brazil (**b**), and Portugal (**c**). Shaded bands are comprised between a more relaxed scenario where declines in incidence are shared in equal parts between cure rate, proportion progressing from primary infection to active disease, and rate of reactivation of latent infection (lower bound) and a more demanding scenario where only reactivation rates are declining (upper bound).

**Extended Data Table 1:**
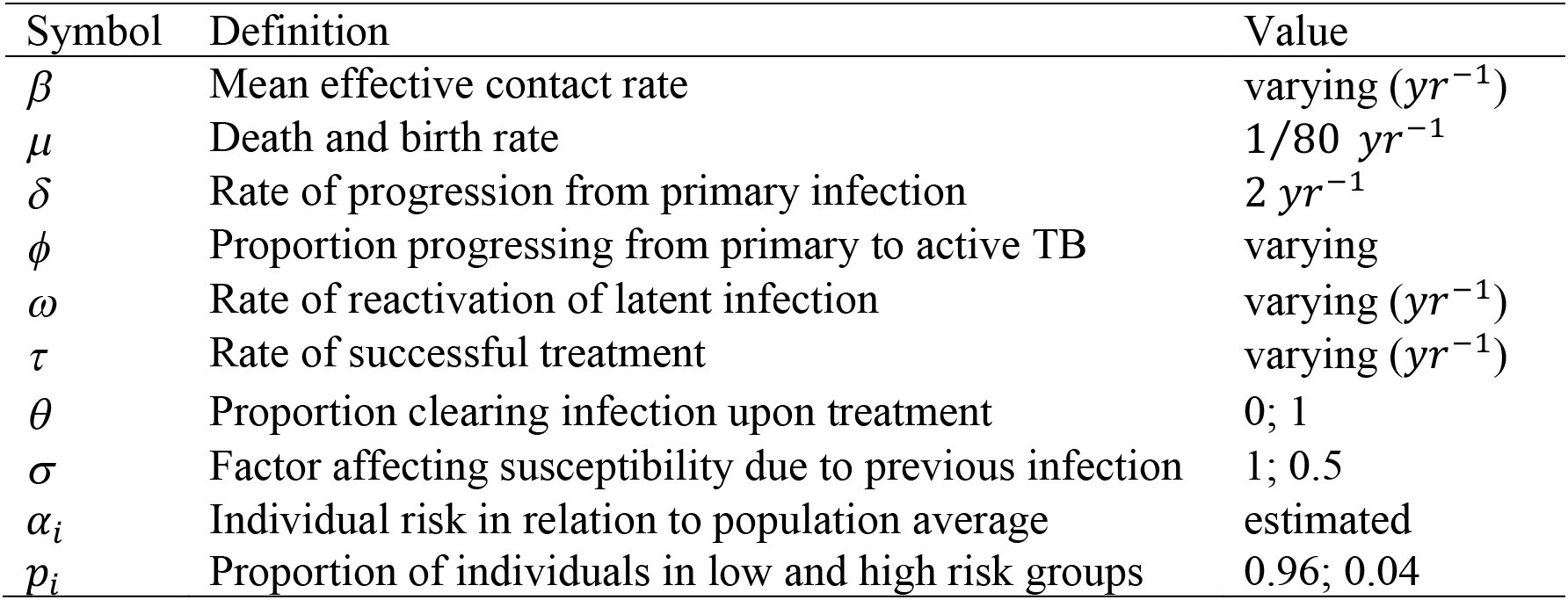
Parameters for tuberculosis transmission model.

**Extended Data Table 2:** Moving target. Values obtained for generating Extended Data Fig. 5.

| Year | Heterogeneous |  | Homogeneous |  |  |  |
| --- | --- | --- | --- | --- | --- | --- |
| | $r_\omega$ | incidence target | $r_\omega$ | | incidence | |
|  |  |  | implemented | perceived | target | achieved |
| 1 | -0.1087 | 49.6 | -0.0732 | -0.0473 | 49.6 | 50.4 |
| 2 | -0.1087 | 47.0 | -0.0765 | -0.0495 | 47.7 | 48.5 |
| 3 | -0.1087 | 44.2 | -0.0804 | -0.0518 | 45.5 | 46.3 |
| 4 | -0.1087 | 41.3 | -0.0852 | -0.0564 | 43.2 | 44.0 |
| 5 | -0.1087 | 38.5 | -0.0909 | -0.0614 | 40.7 | 41.5 |
| 6 | -0.1087 | 35.8 | -0.0980 | -0.0668 | 38.2 | 38.9 |
| 7 | -0.1087 | 33.1 | -0.1076 | -0.0734 | 35.5 | 36.3 |
| 8 | -0.1087 | 30.6 | -0.1220 | -0.0820 | 32.7 | 33.6 |
| 9 | -0.1087 | 28.2 | -0.1479 | -0.0960 | 29.7 | 30.7 |
| 10 | -0.1087 | **26.0** | -0.2150 | -0.1286 | **26.0** | 27.4 |

**Extended Data Table 3:** Parameters for the tuberculosis transmission model utilized to simulate the decline in incidence between 2002 and 2015 in Fig. 2.

| Country | Parameter values |  |
| --- | --- | --- |
|  | Heterogeneous model | Homogeneous model |
| Vietnam | $\beta = 4.59 \text{ yr}^{-1}$<br>$\phi(t) = 0.05e^{-0.0097(t-2002)}$<br>$\omega(t) = 0.0039e^{-0.0089(t-2002)} \text{ yr}^{-1}$<br>$\tau(t) = 2e^{0.0162(t-2002)} \text{ yr}^{-1}$ | $\beta = 10.7 \text{ yr}^{-1}$<br>$\phi(t) = 0.05e^{-0.0245(t-2002)}$<br>$\omega(t) = 0.0039e^{-0.0080(t-2002)} \text{ yr}^{-1}$<br>$\tau(t) = 2e^{0.0363(t-2002)} \text{ yr}^{-1}$ |
| Brazil | $\beta = 3.47 \text{ yr}^{-1}$<br>$\phi(t) = 0.05e^{-0.0031(t-2002)}$<br>$\omega(t) = 0.0013e^{-0.0078(t-2002)} \text{ yr}^{-1}$<br>$\tau(t) = 2e^{0.0041(t-2002)} \text{ yr}^{-1}$ | $\beta = 17.3 \text{ yr}^{-1}$<br>$\phi(t) = 0.05e^{-0.0082(t-2002)}$<br>$\omega(t) = 0.0013e^{-0.0061(t-2002)} \text{ yr}^{-1}$<br>$\tau(t) = 2e^{0.0111(t-2002)} \text{ yr}^{-1}$ |
| Portugal | $\beta = 11.6 \text{ yr}^{-1}$<br>$\phi(t) = 0.05e^{-0.0207(t-2002)}$<br>$\omega(t) = 0.0013e^{-0.0188(t-2002)} \text{ yr}^{-1}$<br>$\tau(t) = 2e^{0.0672(t-2002)} \text{ yr}^{-1}$ | $\beta = 17.1 \text{ yr}^{-1}$<br>$\phi(t) = 0.05e^{-0.0258(t-2002)}$<br>$\omega(t) = 0.0013e^{-0.0185(t-2002)} \text{ yr}^{-1}$<br>$\tau(t) = 2e^{0.0863(t-2002)} \text{ yr}^{-1}$ |

**Extended Data Table 4:**
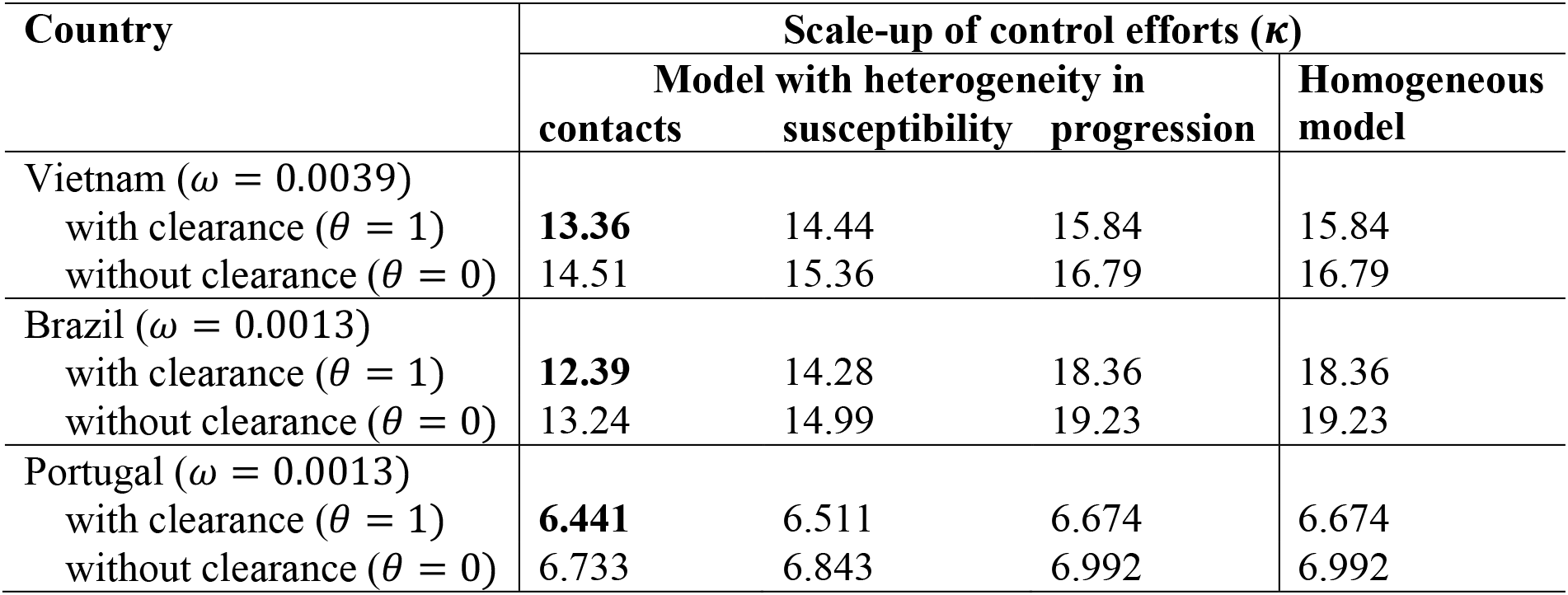
Scale-up of control efforts for meeting End TB incidence target of 90% reduction by 2035. Values corresponding to the model in the Main Text are in bold.

**Extended Data Table 5:** Prevalence of latent TB infection in 2014. Values corresponding to the model in the Main Text are in bold.

| Country | Prevalence of LTBI (%) |  |  |
| --- | --- | --- | --- |
|  | Heterogeneous model | Homogeneous model | WHO region estimate<sup>20</sup> |
| Vietnam ($\omega = 0.0039$) | | | SEA |
| with clearance ($\theta = 1$) | **30.8** | 34.5 | 30.8 [28.3-34.8] |
| without clearance ($\theta = 0$) | 32.7 | 36.9 | |
| Brazil ($\omega = 0.0013$) | | | AMR |
| with clearance ($\theta = 1$) | **16.9** | 22.7 | 11.0 [7.0-20.0] |
| without clearance ($\theta = 0$) | 17.7 | 24.1 | |
| Portugal ($\omega = 0.0013$) | | | EUR |
| with clearance ($\theta = 1$) | **19.7** | 20.3 | 13.7 [9.8-19.8] |
| without clearance ($\theta = 0$) | 20.8 | 21.4 | |

